# Development of optimised human iPSC-derived hepatocytes with improved liver function for *in vitro* metabolic disease modelling and toxicity studies

**DOI:** 10.1101/2024.11.18.624145

**Authors:** Magdalena Łukasiak, Gemma Gatti, Samuel Chung, George Kiloh, Chloe Robinson, Ioannis Kasioulis, Carlos Gil, Lia Panman, Nikolaos Nikolaou

## Abstract

**Background & Aims:** Liver disease is a rising cause of mortality worldwide. Primary human hepatocytes (PHH) and hepatocellular cancer cells are currently used in drug development, however, they come with limitations, including limited supply, rapid loss of function, and tumorigenic origin. In addition, current iPSC differentiation protocols lead to the generation of hepatocyte-like cells with compromised liver-related features. We hypothesised that optimisation of iPSC differentiation protocols can lead to the generation of hepatocyte-like cells with improved metabolic functionality for disease modelling and toxicity screening studies.

**Methods:** Healthy human iPSCs were differentiated to hepatocyte-like cells (Opti-HEP) using a novel 3-step differentiation protocol. Hepatocyte functionality was assessed, including liver maturity marker expression, urea synthesis, *de novo* gluconeogenesis, and CYP450 expression, activity, and induction. Suitability of Opti-HEP to predict drug-induced liver injury (DILI) was evaluated by cell viability assays. CRISPR/Cas9 gene editing was employed to generate *in vitro* inherited metabolic disease models.

**Results:** Opti-HEP expressed similar liver maturity marker levels to those seen in primary human hepatocytes (PHH), in addition to functional urea and gluconeogenesis pathways. CYP450 expression and activity were comparable between Opti-HEP and PHH, with both cell types showing similar levels of CYP3A4 induction upon 1α,25-hydroxy-vitamin D3 treatment. Opti-HEP accurately predicted DILI, following treatment with 7 drugs of known DILI liability. CRISPR-derived Opti-HEP harbouring mutations for inherited metabolic disorders (Ornithine Transcarbamylase Deficiency, Progressive Familial Intrahepatic Cholestasis Type 2, Citrullinemia Type 1) recapitulated key pathophysiological disease features, including reduced protein expression, impaired urea secretion, and bile acid transport.

**Conclusions:** We demonstrate the generation of optimised iPSC-derived hepatocytes with enhanced liver functionality that is comparable to PHH. These data alongside the expansion capacity and amenability of these cells highlight the opportunities this model can offer in the space of disease modelling and large-scale drug efficacy and hepatotoxicity screening.

## 1. Introduction

The liver is one of the most vital organs in the human body performing over 500 critical functions [1]. Its central role in nutrient metabolism, drug detoxification, bile formation, vitamin storage, and immune homeostasis highlights its importance in maintaining overall health. Diseases of the liver, including hepatitis, cirrhosis, fatty liver disease, and hepatocellular carcinoma, are increasing in prevalence and are now a leading cause of morbidity and mortality worldwide [2]. Despite the magnitude of the problem, organ transplant remains the only available option for patients at risk of liver failure, but donor scarcity and the undesired side effects of immunosuppression significantly reduce the likelihood of disease cure. Therefore, there is an urgent need to develop treatments capable of slowing or preventing disease progression.

Accurate liver models are essential in drug discovery to predict human responses to new therapeutic compounds. However, most therapeutics still fail to reach clinical trials or must be withdrawn post-approval due to the lack of translatability between pre-clinical models and the clinic [3,4]. Traditional models, including primary human hepatocytes (PHH) and human hepatocellular carcinoma cell lines have been widely used in drug discovery, but several limitations hinder their effectiveness in drug testing and disease modeling. PHH are considered the gold standard *in vitro* model for liver function studies [5]. However, their supply is limited due to the scarcity of donor livers, and they rapidly decline in culture as they lose their liver-specific functions over time, thus reducing their utility for long-term studies [6]. In addition, significant inter-donor variability amongst different PHH populations affects the reproducibility and predictability of drug response studies during large-scale applications. To overcome some of the issues, a variety of human hepatocellular carcinoma cell lines, including HepG2, HepaRG, Huh7.0 and Hep3b, are often used in liver research, as they remain readily expandable and retain aspects of hepatocyte function. Yet, these cells, being derived from hepatocellular carcinoma, do not fully recapitulate the metabolic and functional profile of primary hepatocytes, lacking many key proteins essential for liver disease modelling [7].

The technology of induced pluripotent stem cells (iPSCs) has offered a promising solution to overcome the limitations of the current liver cell models. iPSCs can be generated from adult somatic cells and differentiated into any cell in the human body, including hepatocytes, providing a renewable source of adult cells. Indeed, iPSCs can proliferate indefinitely and be differentiated towards human hepatocytes on demand, addressing the scarcity of PHH. They express many liver-specific functions, including albumin secretion, glycogen storage, and lipid metabolism, making them more physiologically relevant than hepatocellular carcinoma cell lines [8]. In addition, iPSCs can be derived from patients with specific genetic backgrounds, enabling the study of drug responses and liver diseases in a personalised manner [8]. This is particularly valuable for understanding inter-individual variability in drug metabolism and toxicity. However, and despite their promising potential, iPSC-derived hepatocytes have so far failed to fully replicate the functional capacity and metabolic activity of PHH, due to our lack of understanding of the environmental cues that drive hepatocyte maturation *in vivo* [9]. As a result, most iPSC-derived hepatocytes exhibit immature/foetal-like phenotypes leading to differences in drug metabolism, transporter expression, and response to toxins compared to adult hepatocytes [10,11].

In this study, we decided to generate functional iPSC-derived hepatocytes through direct differentiation following a series of differentiation protocol modifications, including exposure times to certain cytokines, oxygen gradient, and culture media components, aiming to more accurately mimic the molecular mechanisms that occur during liver maturation. Here, we present a comprehensive characterisation of optimised iPSC-derived hepatocytes (Opti-HEP) that demonstrate comparable expression of hepatocyte maturity and transporter markers to those seen in PHH, functional liver metabolic pathways, including Phase I drug metabolism, urea secretion and *de novo* gluconeogenesis, and ability to accurately predict drug-induced liver injury (DILI). Importantly, we demonstrate for the first time, the development of three new CRISPR/Cas9-derived monogenic liver disease models focusing on urea cycle disorders (Ornithine Carbamoyltransferase Deficiency – OTCD, Citrullinemia Type 1 - ASS1) and Progressive Familial Intrahepatic Cholestasis Type 2 (PFIC2), highlighting, in total, the opportunities these cells can offer in the fields of liver disease modeling, drug discovery, and liver toxicity research.

## 2. Materials and Methods

### 2.1. iPSC culture

Ethics for the iPSC lines used in this work were approved under Addenbrooke’s Hospital reference 08/H0311/201. iPSCs were cultured in Vitronectin XF- (#07180; STEMCELL Technologies, Cambridge, UK) coated tissue culture plastic using TeSR-E8 Medium (#05990; STEMCELL Technologies, Cambridge, UK) in a humidified tissue culture incubator at 37°C, 5% CO2. Cell medium was changed daily, and cells were passaged every 5-7 days using ReleSR (#100-0483; STEMCELL Technologies, Cambridge, UK).

### 2.2. HepG2 and PHH culture

HepG2 cells were purchased from LGC Standards (#HB-8065; ATCC, Middlesex, UK) and cultured in Dulbecco’s Modified Eagle Medium (DMEM) (#61870-010; Life Technologies, Waltham, USA) supplemented with 10% FBS (#CS-SA/500; Labtech, East Sussex, UK), 1% penicillin-streptomycin (#P433; Merck Life Science, Dorset, UK) and 1% non-essential amino acids (#11140035; Life Technologies, Waltham, USA) in a humidified tissue culture incubator at 37°C, 5% CO2.

Plateable cryopreserved primary human hepatocytes from three different donors were purchased from CYTES Biotechnologies (#HuHECPMI/6+) and cultured according to manufacturer’s protocol.

### 2.3. iPSC differentiation and Opti-HEP culture

Direct differentiation of human iPSCs towards hepatocytes was performed as previously described [12] with modifications. To initiate differentiation, iPSCs were dissociated using StemPro Accutase Cell Dissociation Reagent (#A1110501; Life Technologies, Waltham, USA) and plated onto gelatine-coated dishes at a density of 20,000 cells/cm^2^. Following completion of differentiation, cells were dissociated using TrypLE (#12604013; Life Technologies, Waltham, USA) and replated onto collagen-coated 96-well plates. For monolayer Opti-HEP culture, cells were kept seeded at a density of 234,000 cells/cm^2^ or 450,000 cells/cm^2^ for transwell culture.

### 2.4. CRISPR/Cas9 gene editing

**OTCD:** To generate the hemizygous single nucleotide mutation D175V in the *OTC* urea cycle gene, a 20-nucleotide PAM-NGG single-guide RNA (sgRNA) was designed to target exon 5. **ASS1:** To generate the homozygous single nucleotide mutation G390R in the *ASS1* urea cycle gene, a 20-nucleotide PAM-NGG sgRNA was designed to target exon 14.

**PFIC2:** To generate the homozygous single nucleotide mutation D482G in the *ABCB11* bile acid transporter gene, a 20-nucleotide PAM-NGG sgRNA was designed to target exon 14.

Repair template single stranded oligodeoxynucleotide (ssODN), containing the desired mutations, were designed with homologous genomic flanking sequences of 50 nucleotides length, centred around the targeted CRISPR/Cas9 cleavage site. Cas9/sgRNA (1/1.5 RNP complex) and 5 μM of single-stranded donor oligonucleotides (ssODN) were delivered to 1×10^6^ single cells by nucleofection using Lonza 4D nucleofection system (#V4XP-3024; Lonza Biologics, Slough, UK) according to the manufacturer’s protocol. Following nucleofection, cells were expanded and seeded in 96-well plates, before DNA was collected, extracted and purified to identify positive clones via Sanger sequencing. The sequences for the sgRNAs, ssODNs, and genotyping primers are illustrated in Suppl. table 1.

### 2.5. Gene expression analysis

Total RNA was extracted using ReliaPrep RNA Cell Miniprep System (#Z6012; Promega, Wisconsin, USA) according to the manufacturer’s protocol. Concentration was determined spectrophotometrically at OD260 on a Nanodrop spectrophotometer (ThermoFisher Scientific, Waltham, MA). Reverse transcription was performed using High-Capacity cDNA Reverse Transcription Kit (#4368813; Applied Biosystems, Waltham, MA).

All quantitative PCR experiments were conducted using an AB StepOne Plus sequence detection system. Reactions were performed in 10l volumes in 96-well plates in reaction buffer containing 5μl TaqMan Fast Advanced Master Mix (#4444557; ThermoFisher Scientific, Waltham, MA). Life Technologies supplied all primers as predesigned Taqman Gene Expression Assays labelled with FAM and endogenous control (*GAPDH, 18S rRNA*, *PPIA*) with VIC. The relative expression ratio was calculated using the ΔΔCt method. Detailed information on the Taqman Gene Expression Assays can be found in Suppl. table 2.

### 2.6. Immunofluorescence

Cells were fixed with 4% p-formaldehyde (#PN28908; ThermoFisher Scientific, Waltham, MA) for 20 minutes, washed three times with DPBS (#D8537; Merck Life Science, Dorset, UK) and blocked for 1 h in 10% donkey serum (#S30-100ML; Merck Millipore, Dorset, UK), 0.1% Triton X-100 (#X100-500ML; Merck Life Science, Dorset, UK) in DPBS. Following blocking cells were subjected to overnight incubation at 4°C with suitable primary antibodies (Suppl. table 3) in 1% donkey serum. Cells were washed three times with DPBS for 5 minutes and incubated for 1 hour with respective secondary antibodies (Suppl. table 3) and DAPI (#10236276001; Merck Life Science, Dorset, UK) for nuclear counterstaining in 1% donkey serum. After washing three times with DPBS for 5 minutes, cells were imaged using a Cellinsight CX7 High Content Analysis Platform (ThermoFisher Scientific, Waltham, MA).

### 2.7. Protein expression and immunoblotting

Total protein was extracted using RIPA lysis buffer (#89900; Life Technologies, Waltham, USA) supplemented with Halt^TM^ protease and phosphatase inhibitor cocktail (#78440; Life Technologies, Waltham, USA). Protein concentrations were measured using the DC^TM^ assay kit (#5000112; Bio-Rad, Hemel Hempstead, UK) according to the manufacturer’s protocol. Gel electrophoresis was run using Criterion gel (#567-8093; Bio-Rad, Hemel Hempstead, UK), and protein samples were transferred onto a polyvinylidene difluoride (PVDF) membrane (#1704157; Bio-Rad, Hemel Hempstead, UK) using the Trans-Blot Turbo Transfer System (#1704150; Bio-Rad, Hemel Hempstead, UK). Membranes were incubated with appropriate primary and secondary antibodies (Suppl. table 4) and normalised to β-actin to ascertain equal gel loading. Bands were visualised with ECL Image Quant LAS 4000 Fluor imager (Amersham, New Jersey, USA), and the protein signal was quantified by densitometry analysis using ImageJ (NIH, Bethesda, MD, http://rsb.info.nih.gov/ij).

### 2.8. CYP450 activity, cell viability, albumin, and urea assays

CYP3A4 activity and induction was measured using P450-Glo CYP3A4 Assay (#V9002; Promega, Wisconsin, USA) according to manufacturer’s protocol. Cell viability was measured using CellTiter-Glo (#G7571; Promega, Wisconsin, USA) according to manufacturer’s protocol. Media albumin levels were measured using Human Albumin SimpleStep ELISA Kit (#ab179887; Abcam, Cambridge, UK), urea levels using QuantiChrom Urea Assay Kit (#DIUR-100; BioAssay Systems, Hayward, CA, USA), and glucose secretion using Glucose Assay Kit (#ab65333; Abcam, Cambridge, UK) according to the manufacturers’ protocols. All data were normalised to total cell number.

### 2.9. Test compound administration

7 DILI-relevant compounds were tested when evaluating DILI predictivity in Opti-HEP and each compound was assigned one of five DILI severity categories according to Proctor et al. 2017 [13] and Williams et al. 2019 [14]. These categories are based on the severity and clinical impact of DILI, as extracted from peer reviewed literature and data contained in product labels ([15]). The criteria used in this classification, the compounds tested in each category and their Cmax are summarised in Suppl. table 5. Compounds in DILI categories 1, 2 and 3 are considered DILI positive and compounds in DILI categories 4 and 5 are considered DILI negative. The compounds were used to dose Opti-HEP in a 9-point dilution where the top concentration of every compound was 250 μM and 0.2% (v/v) DMSO (Dimethyl sulfoxide) concentration. Each compound concentration was tested in triplicate, and data were corrected to maximum viability (0.2% DMSO) and minimum viability controls (250 μM chlorpromazine). All compounds were obtained from the AstraZeneca Compound Library and had their identity and purity confirmed (>98%) with LC-MS.

### 2.10. GalNAc uptake

Synthesis of GalNAc/Cholesterol conjugated siRNAs used was performed by ADViRNA, LLC (Worcester, MA). Duplex RNA oligos (2 nmol) targeting human GAPDH or a non-targeting sequence were conjugated to cholesterol or three N-acetylgalactosamine molecules. A Cy3-label was also linked to the molecule for ease of detection. Preparation and dilution of the duplexes prior the experiment was carried out following manufacturer’s instructions. For the GalNAc uptake, 2 μM of Cholesterol/GalNAc conjugated siRNAs were transfected into Opti-HEP with and incubated for 24 hours, whereupon live cell staining using DAPI (1:10000) was performed. Uptake was measured using a Cellinsight CX7 High Content Analysis Platform (Thermo Scientific, Waltham, MA), and quantified uptake as percentage of counted cells positive for the Cy3 tag signal. For gene expression studies, cells were kept in culture for additional 24 hours, before total RNA was extracted for downstream quantitative PCR analysis.

### 2.11. Bile efflux analysis

For bile acid efflux experiments, 10 µM of Taurocholic Acid (TCA) was added to the lower compartment of both wild-type and ABCB11 mutant transwells for 48 hours. Untreated transwells served as a control. Following 48 hours of treatment, supernatants were collected from both upper and lower compartments of wild-type and ABCB11 mutant transwells, and TCA secretion into the upper transwell compartment was determined by liquid chromatography-Mass-Spectrometry (RapidFire Mass-Spectrometry system; Momentum Biotechnologies, Massachusetts, USA). A deuterated form of TCA (D4-TCA; 100 nM), was used as internal standard. TCA levels were normalised to the total protein.

### 2.12. Statistical analysis

Data are presented as mean ± standard error (SEM), unless stated otherwise. Statistical analysis on gene expression data, CYP3A4 activity, and protein activity was performed using paired or unpaired Student’s t-test. For comparison between different cell types, one-way analysis of variance (ANOVA) followed by Dunnett’s multiple comparison test was employed. Differences were regarded as significant if p-value was < 0.05. Statistical analysis was performed using GraphPad Prism (Graphpad Software Inc., La Jolla, USA).

## 3. Results

### 3.1. Human iPSCs successfully differentiate to hepatocytes, express comparable hepatocyte markers to PHH, and secrete high levels of albumin and urea

Prior to iPSC differentiation, iPSC pluripotency status was investigated by qPCR and immunocytochemistry revealing high expression levels of the gold standard pluripotency markers Octamer-Binding Transcription Factor 4 (*OCT4*), SRY-Box Transcription Factor 2 (*SOX2*), and Nanog Homeobox (*NANOG*) (Fig. 1A-B). Direct differentiation of iPSCs was monitored at different stages until hepatocyte generation, whereupon the characteristic cobblestone morphology of adult hepatocytes was observed, as shown in Fig. 1C. To determine successful differentiation and hepatocyte status, the mRNA levels of *OCT4*, *NANOG*, and *SOX2* were first measured in our Opti-HEP culture, revealing a complete lack of expression, confirming complete differentiation of iPSCs to hepatocytes (Fig. 1A). mRNA and protein expression levels of the gold standard hepatocyte markers albumin (ALB) and hepatocyte nuclear factor 4A (HNF4A) were measured by qPCR and immunocytochemistry, demonstrating comparable mRNA expression levels and % positive cells to those seen in cultures of HepG2 cells and PHH (Fig. 1D-E). The hepatocyte maturity marker alpha-1-antitrypsin (A1AT) was additionally measured in all three cell types revealing comparable percentage levels of A1AT-positive cells between Opti-HEP and PHH, and a complete lack of expression in HepG2 cells. Importantly, our Opti-HEP culture demonstrated low percentage of alpha-fetoprotein-positive cells (11%), and these levels were comparable to those observed in both HepG2 (19%) and PHH (19%) cultures, indicative of the mature phenotype of our cells.

**Figure 1:**
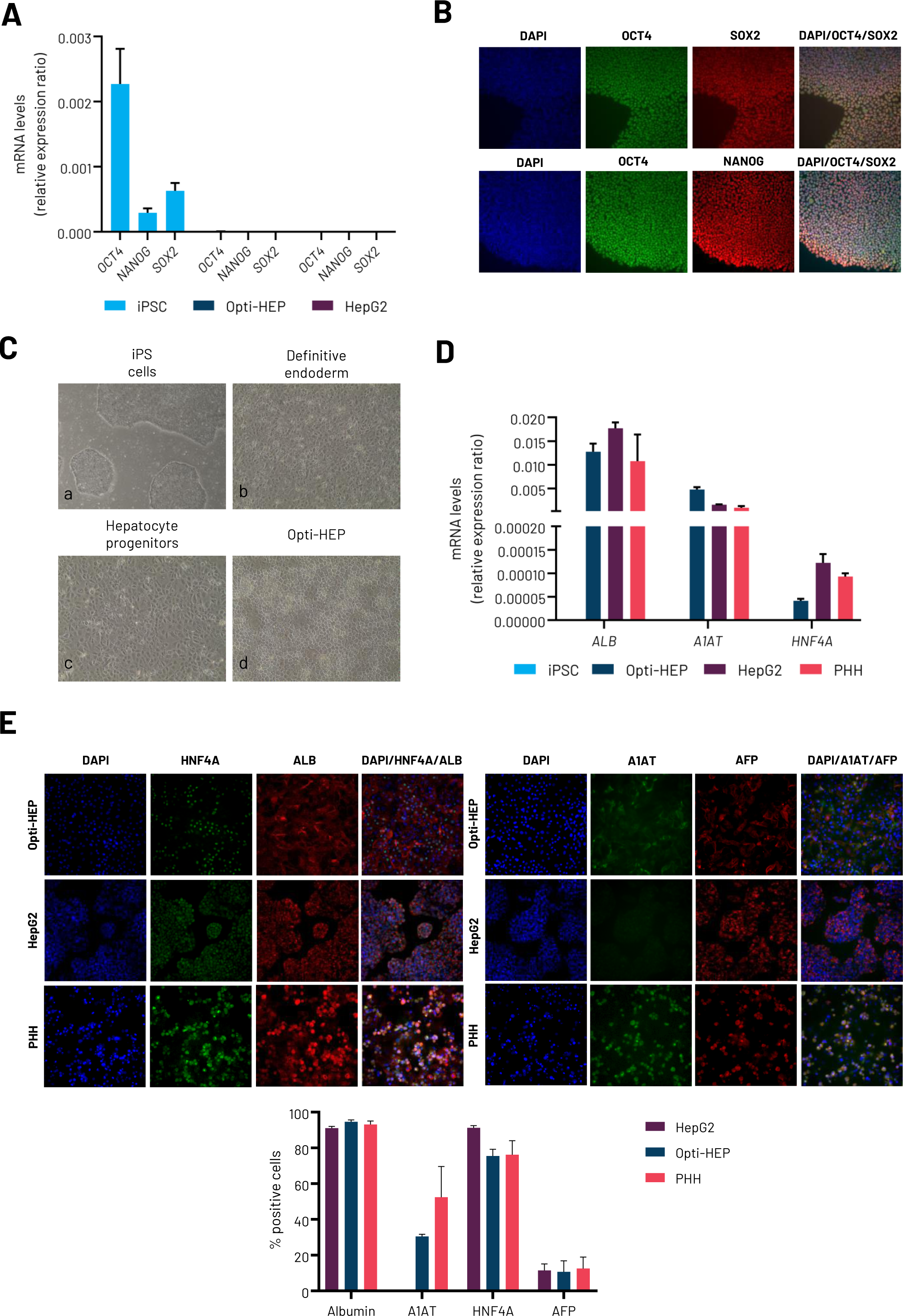
**(A)** mRNA expression levels of Octamer-Binding Transcription Factor 4 (*OCT4*), SRY-Box Transcription Factor 2 (*SOX2*), and Nanog Homeobox (*NANOG*) in wild type induced pluripotent stem cells (iPSCs), wild type Opti-HEP, and HepG2 cells. **(B)** Protein expression of OCT4 (green), SOX2 (red), and NANOG (red) in wild type iPSCs. Objective: 20x: **(C)** Representative brightfield images of key stages of iPSC differentiation towards wild type Opti-HEP. Scale bar: 100 µm. **(D)** mRNA expression levels of key hepatocyte markers albumin (*ALB*), alpha-1-antitrypsin (*A1AT*) and Hepatocyte Nuclear Factor 4 Alpha (*HNF4A*) in iPSCs, Opti-HEP, HepG2 cells, and primary human hepatocytes (PHH). **(E)** Protein expression of albumin (ALB – red), Hepatocyte Nuclear Factor 4 Alpha (HNF4A - green), alpha-1-antitrypsin (A1AT - green) and alpha-fetoprotein (AFP – red) in wild type Opti-HEP, HepG2 cells, and PHH. Objective: 20x RNA data were normalised to housekeeping genes Glyceraldehyde-3-Phosphate Dehydrogenase (*GAPDH*) and Peptidylprolyl Isomerase A (*PPIA*) and are presented as mean±SEM of n=3 independent experiments. Nuclei were counterstained with DAPI (blue). For PHH, cells from three independent donors were used.

Endorsing these findings, mature hepatocyte status and stability of hepatocyte functionality was evaluated by measuring albumin secretion over time. Opti-HEP demonstrated a steady increase of albumin secretion up to day 8-12 post-differentiation, whereupon secretion levels reached PHH albumin secretion levels (Fig. 2A). Crucially, these levels continued to increase until day 24, demonstrating the ability of the Opti-HEP culture to remain functionally stable for a pro-longed period (Fig. 2B).

**Figure 2:**
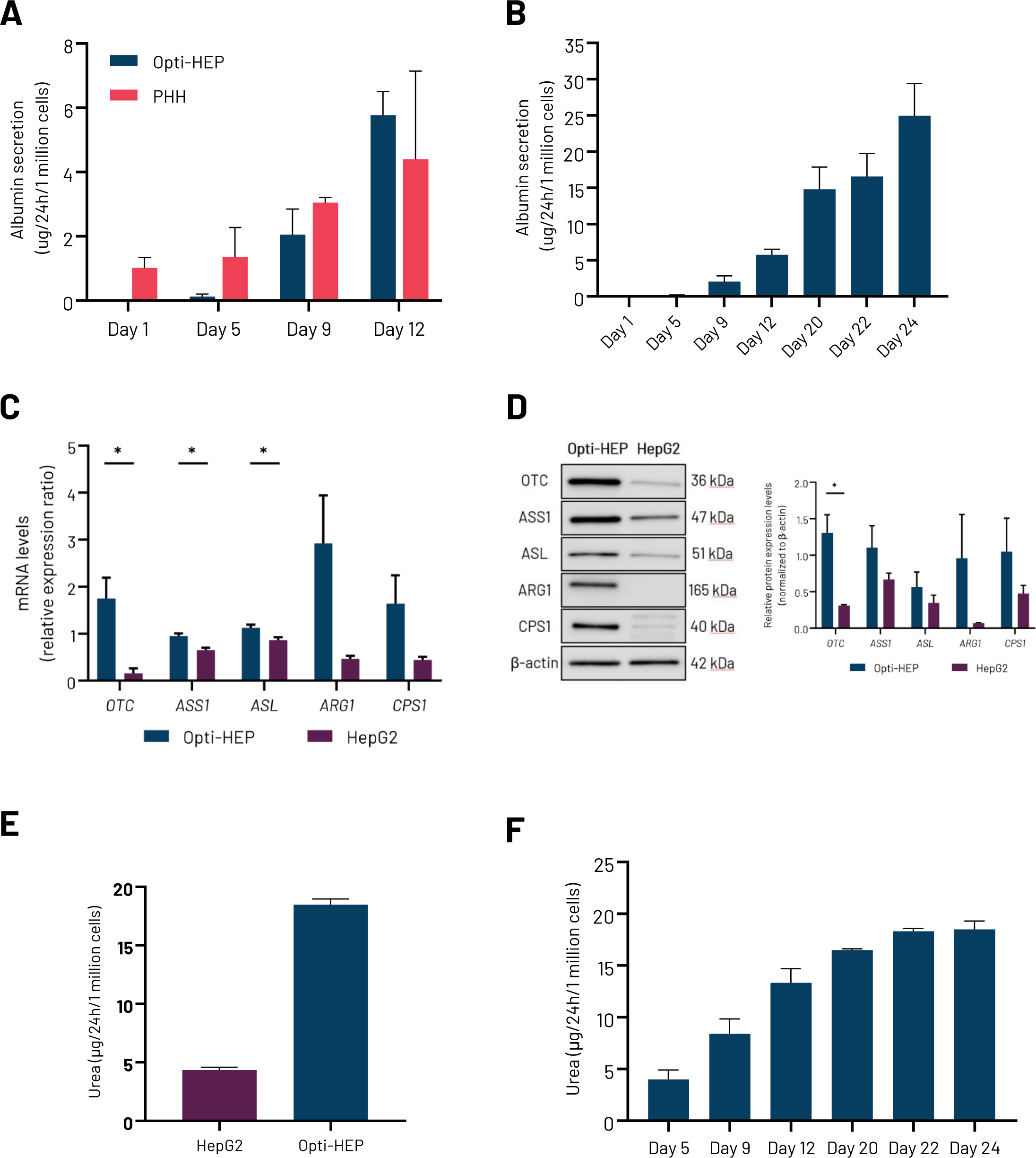
**(A)** Albumin secretion in wild type Opti-HEP and primary human hepatocytes (PHH) over a 12-day period. **(B)** Albumin secretion in wild type Opti-HEP over a 24-day period. **(C)** mRNA expression levels of the urea cycle genes *OTC*, *ASS1*, *ASL*, *ARG1*, and *CPS1* in wild type Opti-HEP and HepG2 cells. **D)** Protein expression levels of the urea cycle enzymes OTC, ASS1, ASL, ARG1, and CPS1 in wild type Opti-HEP and HepG2 cells. Urea secretion by Opti-HEP cells over time. **(E)** Urea secretion in wild type Opti-HEP and HepG2 cells. **(F)** Urea secretion in wild-type Opti-HEP over a 24-day period. mRNA data were normalised to housekeeping gene Peptidylprolyl Isomerase A (*PPIA*). Protein data were normalised to β-actin. Urea secretion data were normalised to cell number. All data are presented as mean±SEM of n=3 independent experiments. For PHH, cells from three independent donors were used. *p<0.05

The liver plays a crucial role in the urea cycle, which is the primary pathway for the detoxification of ammonia in the body. It involves a series of five biochemical reactions that take place both in the mitochondria and the cytoplasm of hepatocytes, converting ammonia and carbon dioxide into urea. To further evaluate the mature hepatocyte status of Opti-HEP, we investigated whether our cells expressed the five main enzymes that are required for a fully functional cycle (Ornithine Transcarbamylase [OTC]; Argininosuccinate synthase 1 [ASS1]; Argininosuccinate lyase [ASL]; Carbamoyl phosphate synthetase I [CPS1]; Arginase 1 [ARG1]). Our results revealed that Opti-HEP expressed significantly higher levels of all five urea cycle enzymes at both mRNA and protein level compared to HepG2 cells (Fig 2C-D). In line with these data, a urea secretion assay demonstrated >3-fold higher media urea levels in Opti-HEP compared to HepG2 cells, and these levels stabilised after approx. 20 days of culture, further confirming the pro-longed stability of our Opti-HEP culture (Fig. 2E-F).

### 3.2. Opti-HEP demonstrate functional *de novo* gluconeogenesis

The liver maintains blood glucose levels during periods of fasting through *de novo* gluconeogenesis, which is the synthesis of glucose from carbohydrate precursors. To confirm the ability of our cells to *de novo* synthesise glucose, we initially investigated whether our cells express glucose-6-phosphatase (*G6PC*), the rate limiting gene of *de novo* gluconeogenesis. Our data revealed higher *G6PC* mRNA levels in Opti-HEP compared to HepG2 cells and closer to those seen in PHH (Fig. 3B). Endorsing these findings, stimulation of *G6PC* expression with the gluconeogenesis inducer dibutyryl-cAMP (dbcAMP; 0.1mM) resulted in >6-fold increase in Opti-HEP, thus demonstrating the inducibility of the pathway in these cells. In contrast, stimulation of HepG2 with dbcAMP resulted in impaired G6PC induction (Fig. 3C). mRNA expression levels of *PCK1* (additional key enzyme in *de novo* gluconeogenesis) were additionally measured and, despite a lower basal expression in Opti-HEP compared to HepG2, induction with dbcAMP was significantly higher with a 10-fold increase in Opti-HEP compared to a 2.5-fold increase seen in HepG2 cells (Suppl. figure 1A-B). Consistent with this data, dbcAMP-induced Opti-HEP demonstrated increased glucose secretion following pyruvate treatment in a dose-dependent manner; however, no response in glucose secretion was observed in HepG2 cells (Fig. 3D).

**Figure 3:**
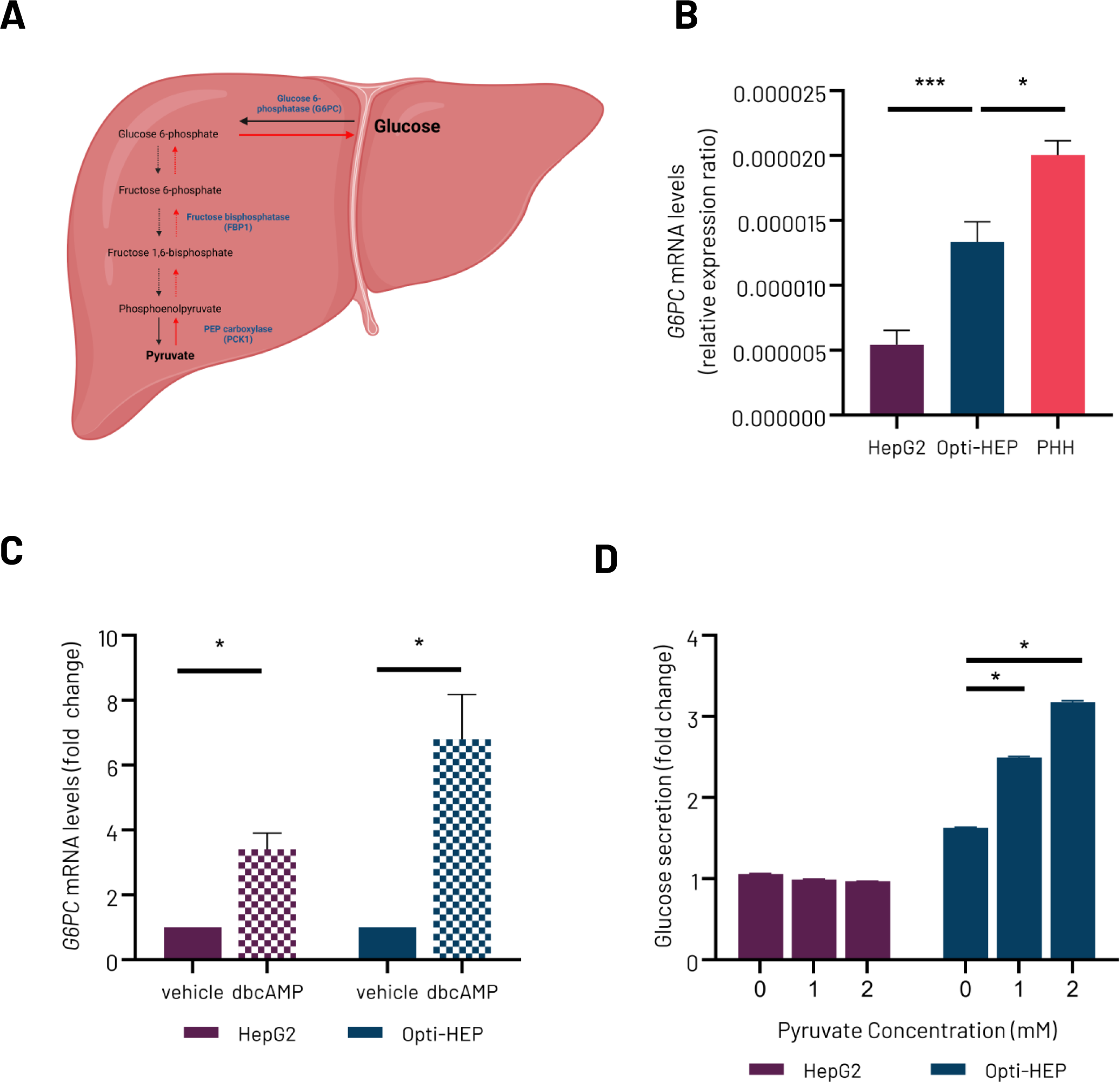
**(A)** Schematic demonstrating the gluconeogenesis pathway present in the liver. Created in BioRender. Nikolaou, N. (2024) https://BioRender.com/b32x291. **(B)** mRNA expression levels of glucose marker glucose-6-phosphatase (*G6PC)* in HepG2 cells, wild type Opti-HEP, and primary human hepatocytes (PHH). **(C)** mRNA expression levels of *G6PC* following stimulation with the gluconeogenesis inducer dibutyryl-cAMP (dbcAMP; 0.1 mM) in HepG2 cells and wild type Opti-HEP. Data presented as mean+SEM of 3-4 independent experiments. **(D)** Glucose secretion in vehicle and dbcAMP treated HepG2 cells and wild type Opti-HEP upon pyruvate challenge. mRNA expression data were normalised to housekeeping gene *18SrRNA*. Glucose secretion data were normalised to cell number. All data presented as mean+SEM of 3-4 independent experiments. *p<0.05; ***p<0.001.

### 3.3. Opti-HEP express ASGR1 receptor and show proper protein membrane localisation and function

To investigate whether Opti-HEP can be used as a potential platform for GalNAc-conjugated therapeutics, we initially determined the expression levels of asialoglycoprotein receptor 1 (ASGR1) as well as its cellular localisation and functional activity in our cells. *ASGR1* mRNA expression was measured in undifferentiated iPSCs, HepG2, Opti-HEP, and PHH revealing comparable levels between Opti-HEP and PHH; iPSCs did not show any *ASGR1* expression, whilst HepG2 showed the highest expression amongst all cell types (Fig. 4A). Despite the high expression levels, HepG2 cells failed to demonstrate proper cellular localisation of ASGR1 protein in the hepatocyte membrane as shown by immunocytochemistry. In contrast, ASGR1 staining in Opti-HEP revealed the receptor expression in endomembrane compartments as well as proper localisation in the basolateral membrane, suggesting ability of the receptor to exploit its internalisation function (Fig. 4B). Indeed, proof-of-concept uptake experiments in Opti-HEP showed approx. 60% Cy3 fluorescent tag-positive Opti-HEP following transfection with a Cy3-tagged GalNAc-siRNA conjugate targeting GAPDH; in contrast, only 10% of human iPSC were found positive for Cy3, supporting the specific role of ASGR1 in the internalisation of this class of therapeutics (Fig. 4C). Finally, mRNA expression analysis revealed a 50% reduction in *GAPDH* mRNA levels in Opti-HEP – but no response in iPSCs – confirming, in total, the presence of a fully functional ASGR1 in our cells (Fig. 4D).

**Figure 4:**
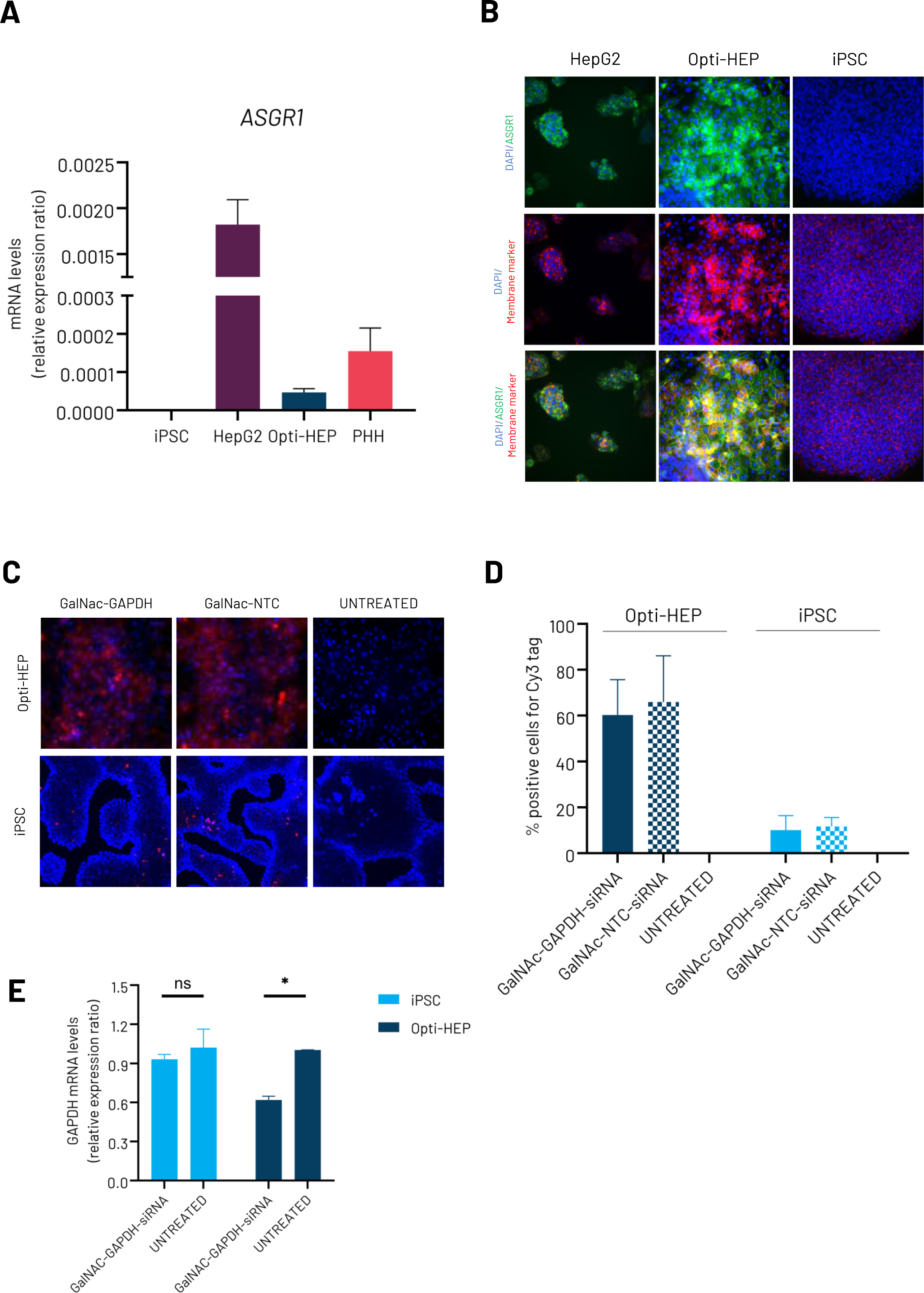
**(A)** mRNA expression levels of asialoglycoprotein receptor 1 (*ASGR1*) in wild type induced pluripotent stem cells (iPSC), HepG2 cells, wild type Opti-HEP, and primary human hepatocytes (PHH), HepG2 and iPSC; **(B)** Representative immunocytochemistry pictures demonstrating staining of the ASGR1 subunit protein in HepG2 cells, wild type Opti-HEP and iPSC. Objective: 20x. **(C)** Representative immunocytochemistry pictures demonstrating uptake (red) of 2 μM GalNAc-siRNA conjugates targeting GAPDH following 24 hours of treatment in wild type Opti-HEP and iPSC. GalNAc-NTC conjugate was used as positive control. Objective: 20x. **(D)** Quantification of uptake expressed as % of positive cells for Cy3 staining in Gal-NAc-GAPDH-siRNA treated-, Gal-NAc-NTC-siRNA treated-, or untreated iPSC and wild type Opti-HEP. **(E)** mRNA expression levels of *GAPDH* in iPSC and Opti-HEP following 48 hours of treatment with 2 μM Gal-NAc-GAPDH-siRNA compared to untreated cells. mRNA levels were normalised to housekeeping genes *PPIA* and *18SrRNA*. Nuclei were counterstained with DAPI and membrane markers with Phalloidin/E-cadherin. All Opti-HEP data are presented as mean±SD of n=3 independent experiments. iPSC data are presented as mean±SD of n=1 experiment.

### 3.4. Opti-HEP demonstrate functional cytochrome P450 expression and activity

To confirm whether iPSC-derived Opti-HEP exhibit drug-metabolizing potential, we checked the expression profile of the Phase I CYP450 enzymes in our cells, showing comparable expression levels of *CYP3A4*, *CYP2B6*, *CYP2C9*, *CYP2C19* and *CYP2J2* between Opti-HEP and PHH, and significantly higher to those seen in HepG2 cells. Interestingly, high expression levels of *CYP2A6* were also measured in our cells, but no expression in PHH was detected (Fig. 5A). Additional analysis on the expression of *CYP1A2* and *CYP2E1* was performed, and mRNA levels were found similar to those seen in HepG2 cells (Suppl. figure 2A-B). Consistent with the high *CYP3A4* mRNA levels, Opti-HEP treatment with rifampicin (50 μM) and 1α,25-hydroxy-vitamin D3 (100 nM) resulted in >6- and >17-fold increase in *CYP3A4* mRNA expression compared to vehicle-treated cells, respectively, whilst, measurement of basal CYP3A4 activity using P450-Glo™ assays revealed comparable activity between Opti-HEP and PHH and higher to that seen in HepG2 cells (Fig. 5B-C). In line with Fig. 5B, treatment of all three cell lines with increasing concentrations of 1α,25-hydroxy-vitamin D3 (10-100 nM) resulted in approx. 3-fold induction of CYP3A4 activity in both Opti-HEP and PHH, whilst no induction was observed in HepG2 cells. Additionally, co-treatment of all cell types with ketoconazole, a potent CYP3A4 inhibitor, was able to inhibit the CYP3A4 activity, demonstrating, in total, the ability of our cells to be used for both CYP3A4 induction and inhibition studies.

**Figure 5:**
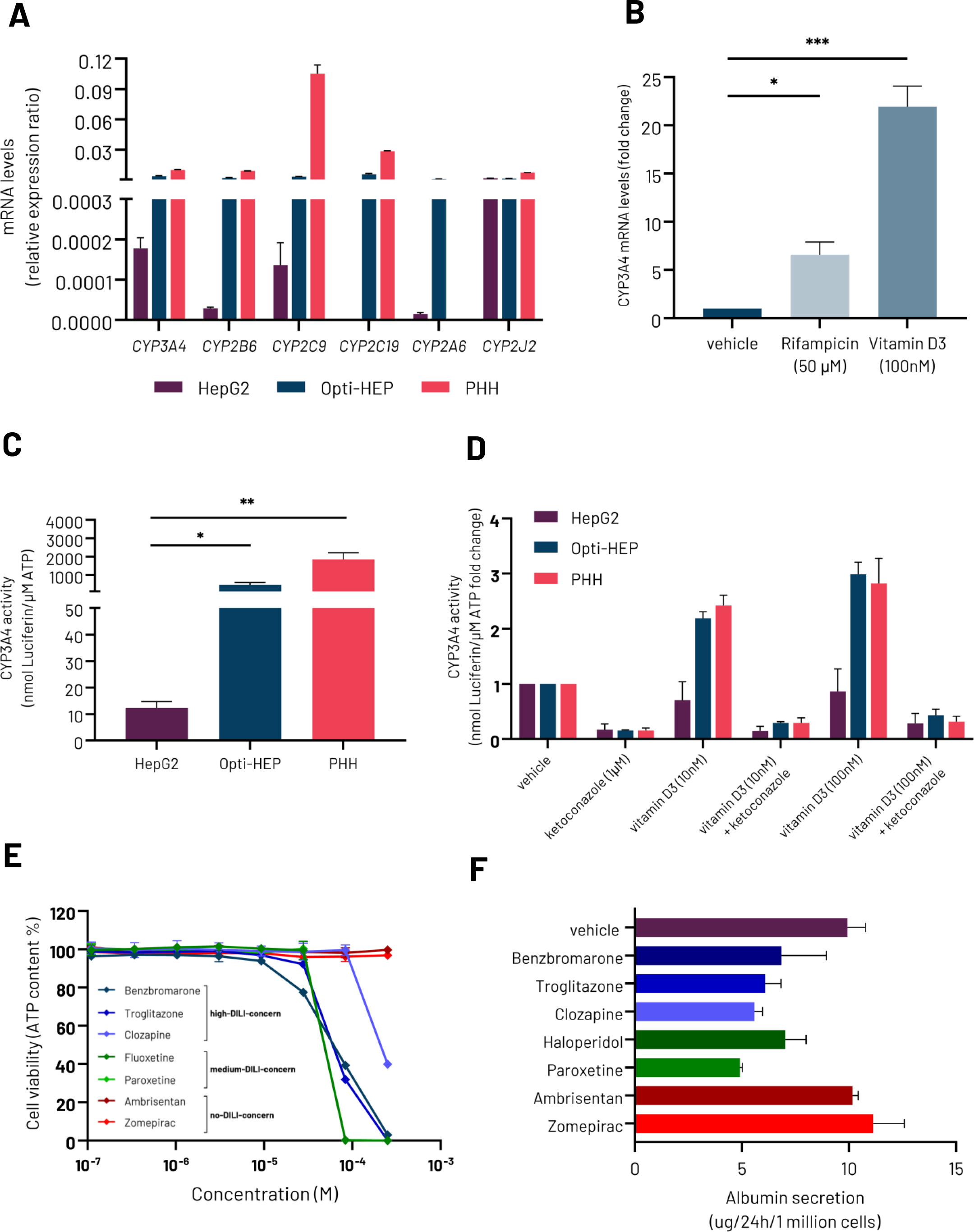
**(A)** mRNA expression levels of Phase I CYP450 genes in HepG2 cells, wild-type Opti-HEP, and primary human hepatocytes (PHH). **(B)** *CYP3A4* mRNA induction in Opti-HEP following 72 hours treatment with vehicle (DMSO), 50 μM Rifampicin, or 100 nM vitamin D3 (CYP3A4 inducers). **(C)** Basal CYP3A4 activity in HepG2 cells, wild type Opti-HEP, and PHH. **(D)** CYP3A4 induction and inhibition in HepG2 cells, wild type Opti-HEP, and PHH, following 72 hours of treatment with vehicle (DMSO), 10 nM or 100 nM vitamin D3 alone or a combination with 1 μM ketoconazole (CYP3A4 inhibitor, 1 h incubation). **(E)** Cell viability (ATP content) in Opti-HEP following 48 hours of treatment with increasing concentrations (0-250 μM) of 7 compounds with known DILI liability (classification based on Proctor *et al*. 2017 [13]). **(F)** Albumin secretion in Opti-HEP following 48 hours of treatment with 250 μM of 7 compounds with known DILI liability (classification based on Proctor *et al*. 2017 [13]). mRNA data were normalised to housekeeping gene *18SrRNA* and are presented as mean±SEM of n=3-4 independent experiments. CYP3A4 activity data were normalised to ATP levels and are presented as mean±SEM of n=3-5 independent experiments. Cell viability data were normalised to positive (250 μM chlorpromazine) and negative controls (0.2% DMSO) and are presented as mean±SD of n=3 technical replicates. For PHH data, cells from 3 independent donors were used. *p<0.05; **p<0.01; ***p<0.001.

Finally, to determine whether Opti-HEP can be used for Drug-Induced Liver Injury (DILI) predictivity studies, we additionally screened our cells against a set of 7 DILI-relevant compounds using cytotoxicity (ATP) and albumin secretion as endpoint assays. Both assays revealed the ability of Opti-HEP to accurately predict DILI, with the high- and medium-DILI-concern related compounds demonstrating increased cytotoxicity and decreased albumin secretion, whilst the no-DILI-concern related drugs had no effect in either cell viability or metabolic output (Fig. 5E-F).

### 3.5. Opti-HEP can be used to model urea cycle disorders *in vitro*

Urea cycle disorders (UCDs) represent a group of metabolic disorders characterised by deficiency in any of the enzymes within the urea cycle, and are associated with a range of life-threatening symptoms, including developmental delay, cerebral oedema, and coma. Despite their severity, there are currently no licensed treatments for any of the UCDs, mainly due to the lack of predictive *in vitro* liver models for drug efficacy screening studies. Following confirmation of urea cycle functionality in Opti-HEP cells, we investigated whether those cells could be a suitable platform for modelling urea cycle disorders. We aimed to develop two iPSC-derived UCD models, focusing on Ornithine Transcarbamylase deficiency (OTCD) (the most common urea cycle disorder [UCD] with an estimate disease incidence of 1:60,000-70,000) and Citrullinemia Type 1 (ASS1 deficiency; estimated disease incidence of 1:250,000).

To this end, we introduced two disease-causing mutations, either the missense mutation D175V (Asp175Val) in the *OTC* gene or the missense G390R (Gly390Arg) mutation in *ASS1* gene using CRISPR/Cas9. Successful introduction of the mutations was confirmed by Sanger sequencing (Fig. 6A-B), with both mutant iPSC lines retaining their pluripotency status, as shown by qPCR and ICC (Suppl. figure 3A-C). Complementing these data, both iPSC lines successfully differentiated towards Opti-HEP, and mRNA expression analysis revealed no differences in the gene expression levels of *ALB*, *A1AT*, and *HNF4A* compared to their isogenic controls (Fig.6C-D).

**Figure 6:**
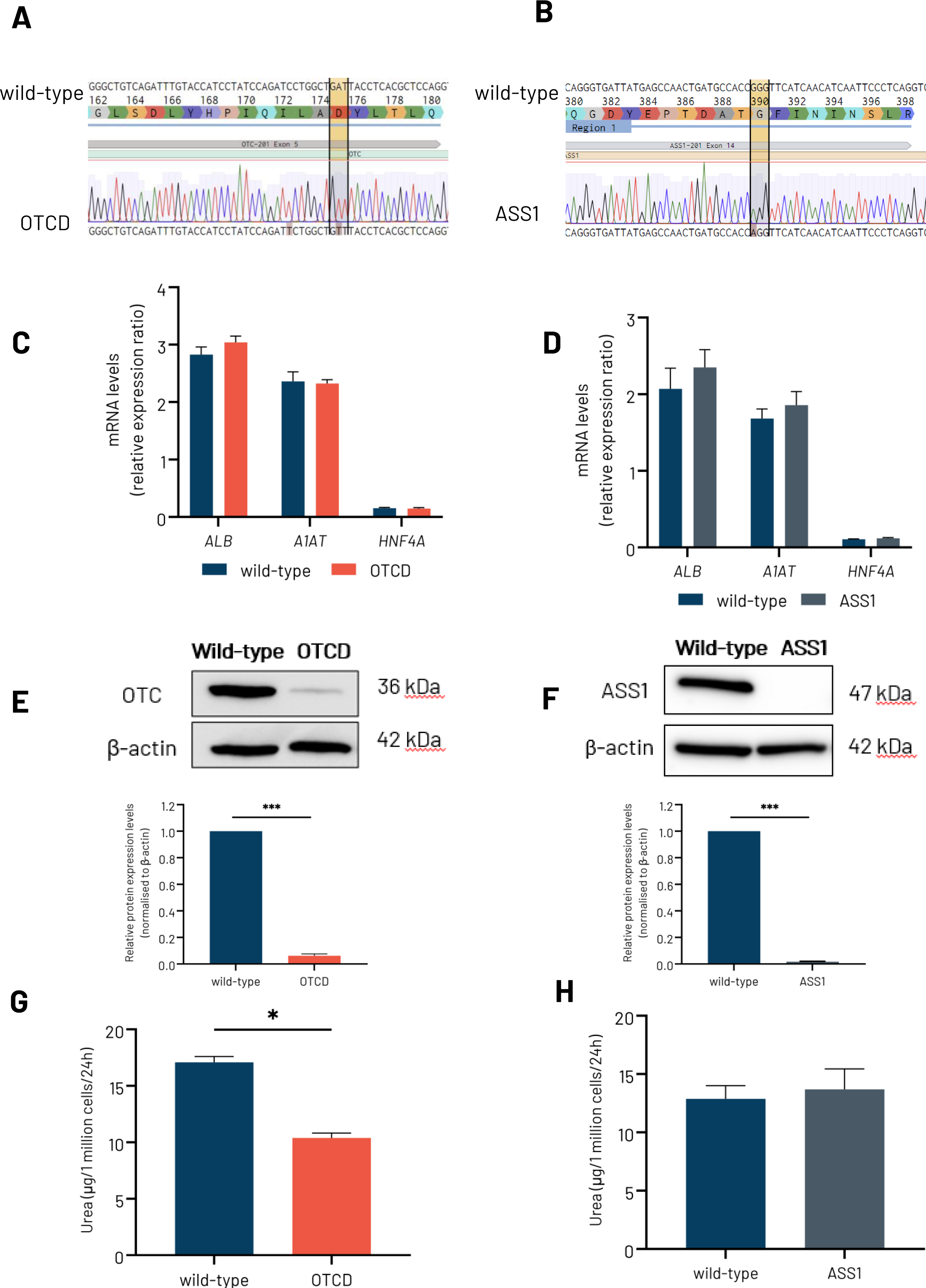
**(A)** Sanger sequencing chromatogram showing wild type and mutated induced pluripotent stem cells (iPSC) carrying the D175V mutation (GAT>GTT) in the OTC gene. The codon change is highlighted with yellow. **(B)** Sanger sequencing showing healthy wild type and mutated iPSCs carrying the G390R mutation (GGG>AGG) in the ASS1 gene. The codon change is highlighted with yellow. **(C)** mRNA expression levels of the key hepatocyte markers albumin (*ALB*), alpha-1-antitrpysin (*A1AT*), and Hepatocyte Nuclear Factor 4A (*HNF4A*) in wild type and ornithine transcarbamylase deficiency (OTCD) Opti-HEP. **(D)** mRNA expression levels of the key hepatocyte markers albumin (*ALB*), alpha-1-antitrpysin (*A1AT*), and Hepatocyte Nuclear Factor 4A (*HNF4A*) in wild type and Citrullinemia type I (ASS1) Opti-HEP. **(E)** Protein expression of ornithine transcarbamylase (OTC) in wild-type and OTCD Opti-HEP. **(F)** Protein expression of argininosuccinate synthetase 1 (ASS1) in wild-type and ASS1 Opti-HEP. **(G-H)** Urea secretion levels in wild-type and OTCD and ASS1 Opti-HEP. mRNA data were normalised to housekeeping gene *18SrRNA* and protein data to β-actin. Urea secretion data were normalised to total cell number. All data are presented as mean±SEM of n=2-3 independent experiments. *p<0.05; ***p<0.001.

Confirming the detrimental role of the OTC and ASS1 mutations on protein stability, western blotting analysis revealed >90% reduction in OTC and ASS1 protein expression in the OTC- and ASS1-mutant Opti-HEP, respectively (Fig. 6E-F). Finally, in line with the protein data, urea secretion was decreased by 40% in OTC-mutant Opti-HEP compared to wild-type; however, no differences in urea secretion were observed in the ASS1-mutant cells (Fig. 6G-H).

### 3.6. Opti-HEP can be used to model Bile Salt Export Pump Deficiency *in vitro*

In addition to modelling UCDs *in vitro*, we further investigated whether our cells could offer an efficient platform to mimic liver monogenic diseases, for which functional *in vitro* human disease models are limited. We focused on the development of a human bile acid transport cell model characterised by deficiency in the bile salt export pump (ABCB11/BSEP). Defects in BSEP have been associated with a range of clinical phenotypes, including familial intrahepatic cholestasis type 2 (PFIC2), benign recurrent intrahepatic cholestasis type 2, and intrahepatic cholestasis of pregnancy [16,17]. Amongst these, PFIC2 is the most severe form, however, there are currently no effective licenced treatments due to the lack of efficient human disease models to study the molecular mechanisms underpinning disease phenotype.

To initially confirm that our cells can be used for the development of an *in vitro* BSEP deficiency model, we measured the gene expression levels of *ABCB11* - as well as the additional efflux transporters *ABCG2* and *ABCC2* - in HepG2 cells, Opti-HEP, and PHH, revealing comparable expression between Opti-HEP and PHH (Fig. 7A). Following confirmation of adequate *ABCB11* expression, a CRISP/Cas9-derived ABCB11-mutant iPSC line focusing on the common D482G missense mutation was generated, and the single point mutation GAT > GGT was introduced in exon 14 of the *ABCB11* gene (Suppl. figure 4A). Presence of the D482G mutation in *ABCB11* was confirmed by Sanger sequencing and, similar to the OTC and ASS1 clones, the ABCB11 D482G mutant iPSCs showed high levels of expression of pluripotency markers OCT4, SOX2, and NANOG as measured by immunocytochemistry (Suppl. figure 4B-C).

**Figure 7:**
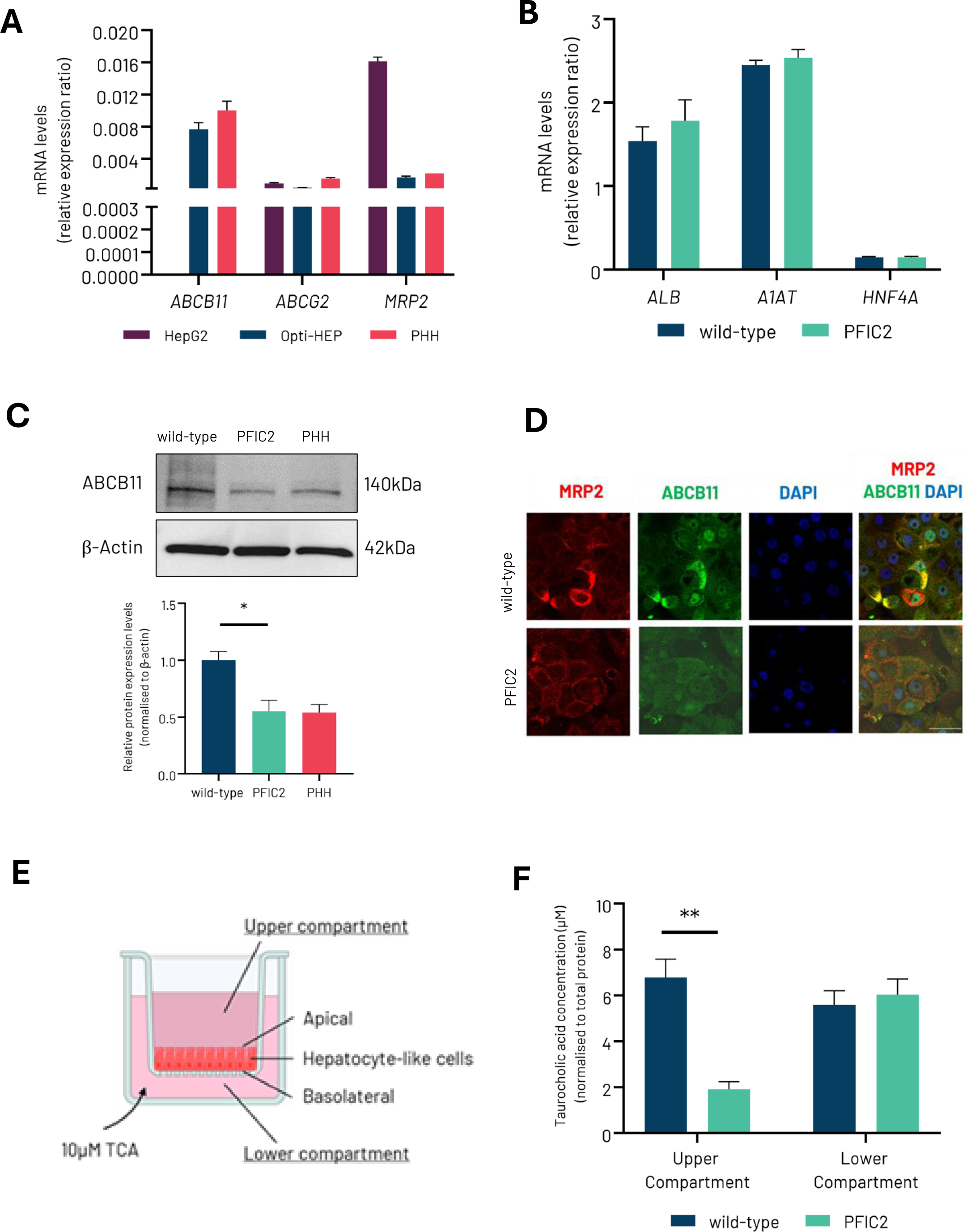
**(A)** mRNA expression levels of the hepatocyte transport markers *ABCB11*, *ABCG2* and *ABCC2* in HepG2 cells, wild type Opti-HEP, and primary human hepatocytes (PHH). **(B)** mRNA expression levels of the key hepatocyte markers albumin (*ALB*), alpha-1-antitrpysin (*A1AT*), and Hepatocyte Nuclear Factor 4A (*HNF4A*) in wild type and progressive familial intrahepatic cholestasis type 2 (PFIC2) Opti-HEP. **(C)** Protein expression levels of ABCB11 in wild type Opti-HEP, PFIC2 Opti-HEP and primary human hepatocytes (PHH). **(D)** Representative immunocytochemistry pictures showing MRP2 and ABCB11 expression and localisation in wild type and PFIC2 Opti-HEP when cultured in transwells. Objective: 20x. **(E)** Schematic depiction of the polarisation of Opti-HEP in transwells, and the addition of 10 µM taurocholic acid (TCA) to the lower compartment for 48 hours. **(F)** TCA levels into the upper and lower transwell compartment of wild type and PFIC2 Opti-HEP following 48 hours of treatment with 10 µM TCA initially added to the lower compartment. mRNA data were normalised to housekeeping gene *18SrRNA* and protein data to β-actin. TCA secretion data were normalised to total protein. Nuclei were counterstained with DAPI. All data are presented as mean±SEM of n=3-4 independent experiments. *p<0.05; **p<0.01.

To achieve hepatocyte polarisation, both wild-type and ABCB11-mutant iPSCs were differentiated into Opti-HEP and, upon completion of differentiation, cultured in transwells. Transwell culture increased, as expected, basal *ABCB11* expression compared to monolayer cultures of Opti-HEP (Suppl. Figure 4D). mRNA expression analysis of the hepatocyte maturity markers *ALB*, *A1AT*, and *HNF4A* revealed no differences between wild-type and ABCB11-mutant (PFIC2) Opti-HEP, confirming successful differentiation of both cell lines towards hepatocytes (Fig. 7B). Western blot analysis in both cell lines demonstrated reduced ABCB11 protein levels in PFIC2 Opti-HEP compared to isogenic control and this effect was accompanied by a complete lack of ABCB11 expression in the Opti-HEP membrane, as measured by immunocytochemistry (Fig. 7C-D). No significant differences in MRP2 localisation between wild-type and PFIC2 Opti-HEP were observed (Fig. 7D). Finally, bile acid efflux was studied following 48 hours of incubation with 10 µM taurocholic acid (TCA), as shown in Fig. 7E. Confirming the lack of functional ABCB11 protein in PFIC2 Opti-HEP, a 3-fold reduction in TCA secretion into the upper transwell compartment of the mutant cells was observed when compared to wild-type Opti-HEP, highlighting impaired bile acid efflux in our disease model (Fig. 7F).

## 4. Discussion

In this study, we have presented a comprehensive characterisation of optimised iPSC-derived hepatocytes, demonstrating their superior functionality compared to HepG2 cells that is, in many aspects, comparable to primary human hepatocytes. Our *in vitro* experiments have shown high and stable expression of hepatocyte maturity markers and transporters, as well as functional urea, *de novo* gluconeogenesis, and Phase I metabolism pathways, highlighting their ability to be used as novel models for large-scale drug efficacy and toxicity screening studies. Importantly, we have demonstrated their ability for use in disease modelling by developing three novel monogenic liver disease models focusing on UCDs and PFIC2.

Over the past fifteen years, a multitude of iPSC differentiation protocols have been developed focusing on the generation of functional iPSC-derived hepatocytes [17–21]. The majority of these protocols mimic the stages seen during liver development *in vivo* by using a variety of soluble factors, aiming to recapitulate the molecular and cellular cues observed in human liver [18–22]. However, most of them fail to generate fully maturated hepatocytes, especially with respect to urea and drug metabolism-related pathways, thus rendering their use for metabolic and drug toxicity studies not possible [11,23,24]. In our study, we have shown that, unlike HepG2 cells, our cells have a fully functional urea cycle pathway as all key urea cycle enzymes are expressed at mRNA/protein levels consistent with urea secretion. This agrees with previous data showing impaired urea cycle in HepG2 cells, an effect that is possibly related to the lack of OTC and ARG1 expression in this cell line [25]. In addition to urea, our cells have revealed fully functional Phase I metabolism, as evident by high expression and activity levels of the P450 enzymes as well as ability to accurately predict liver injury. These data alongside our additional work on the characterisation of glucose metabolism within Opti-HEP underscore the metabolic competency of our cells.

Recently, alternative methodologies relying on the over-expression of hepatocyte nuclear factors combined with different nuclear receptors have been developed, aiming to generate functional iPSC-derived hepatocytes through a more simplified process [9]. The authors have described the generation of iPSC-derived hepatocytes that display functional features of primary human hepatocytes, including liver maturity marker, Phase I/II, and carbohydrate/lipid marker expression alongside basal CYP3A4 activity. However, these cells show evidence of functionality decline following prolonged culture, in addition to lack of characterisation against major liver metabolic pathways, including urea secretion, functional *de novo* gluconeogenesis, or CYP3A4 induction/inhibition.

The asialoglycoprotein receptor 1 (ASGR1) is known to be expressed within a subpopulation of functional hepatocytes, therefore representing an important marker for mature and homogeneous iPSC-derived hepatocyte cultures. In this regard, Peters et al. [26] have shown ASGR1-positive iPSC-derived hepatocytes share a transcriptional profile that is more similar to adult hepatocytes than ASGR1-negative cells, accompanied by high expression levels of genes involved in synthetic function (e.g., albumin, alpha-1 antitrypsin), energy metabolism (e.g., HNF4A), bile production/metabolism, and drug detoxification pathways. Opti-HEP were able to recapitulate the expression of mature hepatocyte markers, including ASGR1, as well as several hepatic functions similar to primary human hepatocytes, thus confirming the robustness of the differentiation protocol and maturity status of our cells.

Recent developments in drug discovery and development areas have focused on the use of RNA-based therapies, which represent an attractive solution to modulate clinically relevant targets that are considered unattackable via traditional approaches. *In vivo* liver delivery of RNA-based therapies can be achieved by conjugating the RNA molecule to a carbohydrate moiety, N-acetylgalactosamine (GalNAc), that facilitates receptor-mediated uptake of the molecule. In this regard, the abundant and specific hepatic ASGR1 is capable of mediating uptake and internalisation of GalNAc-conjugated molecules allowing this class of therapeutics to overcome the struggle of cellular barriers [27]. Previous studies evaluating antisense oligonucleotide (ASO) internalisation and related knockdown activity towards a specific target using Huh7 and HepG2 cells showed no to little correlation between GalNAc uptake and knockdown activity, highlighting the limitations of liver carcinoma cell lines in GalNAc-mediated screening [28]. In line with this study, our ICC data revealed ASGR1 expression in HepG2 cells, but lack of proper transporter localisation; in contrast, Opti-HEP overcome the limitations of liver carcinoma cells as evident by ASGR1 membrane localisation and reduced GAPDH levels following GalNAc-siRNAGAPDH treatment.

Modelling inherited monogenic liver diseases *in vitro* using traditional hepatic cell models has been a long-standing challenge, predominantly due to the limitations of hepatocellular cell lines and PHH in expressing the necessary metabolic pathways and controlling donor variability, respectively. The development of iPSC-derived hepatocytes has provided an alternative and effective strategy, allowing researchers to study disease mechanisms and screen novel therapeutics *in vitro* in addition to investigating population diversity. Given the liver-specific expression of the urea cycle pathway, patient-derived iPSCs have been previously utilised to generate disease hepatocytes for the study of UCDs; indeed, a few studies have demonstrated the generation of functional hepatocytes from patients with mutations in all urea cycle genes [29]. In this study, we have demonstrated, for the first time, the development of two genetically engineered UCD hepatocyte models, OTCD and Citrullinemia type 1, revealing significant decreases in protein expression alongside impaired urea secretion. OTCD is an X-linked genetic disease and the most common UCD accounting for almost half of all urea cycle defects [30]. To date, >400 mutations, ranging from amino acid replacements to RNA splicing defects, premature protein terminations and deletions, have been reported, however, there is no prevalence, and all mutations may equally arise in any of the 10 exons of the OTC gene [31,32]. In our work, we focused on the D175V mutation (Asp175Val), a deleterious mutation first identified in a female patient in 1997 that was characterised by deficient enzyme activity accompanied by a range of neurological symptoms, including episodes of confusion, disorientation, and altered environmental contact [32]. Supporting the clinical symptoms, our data reveal, for the first time, significant decreases in OTC protein levels in D175V-mutant cells in addition to lower urea secretion levels. In this regard, our data are in line with previous studies, where differentiation of patient-derived iPSCs harbouring different mutations in the OTC gene resulted in decreased OTC protein levels and urea secretion compared to healthy controls [33,34]. Similar to OTCD, Citrullinemia Type I is a UCD caused by defects in the ASS1 gene, and encompasses a variety of clinical phenotypes, including hyperammonemia, elevated plasma citrulline and low plasma arginine, as well as elevated urinary orotic acid levels [35]. At least 137 mutations in the ASS1 gene have been described, with the mutation G390R (Gly390Arg) being the most frequent and globally spread throughout the population [35,36]. Previous endeavours to generate iPSC-derived hepatocytes from patient iPSCs harbouring either the G390R or additional disease-related mutations (G259*; R304W) resulted in functional hepatocytes but failed to fully recapitulate the disease phenotype. Similarly, in our CRISPR/Cas9-derived model, whilst we demonstrated significant differences in ASS1 expression between G390R Opti-HEP and isogenic controls, we did not observe any changes in urea secretion, reflecting the complexity of the metabolic pathways related to ASS1 activity. Indeed, in addition to the metabolism of citrulline towards argininosuccinate formation to feed the urea cycle, ASS1 plays a central role in the recycling of nitric oxide through the citrulline-nitric oxide cycle, thus highlighting the need to establish additional phenotypic assays to properly characterise these *in vitro* cell models [37].

PFIC2 affects approx. 1 in 50000 to 100000 births worldwide and is associated with a variety of clinical features, ranging from giant cell transformation, inflammation, and canalicular cholestasis at neonatal stage, to canalicular cholestasis and fibrosis at later stages in life [38,39]. At least 200 different mutations in the ABCB11 have been reported, with the most common ones being the p.E297G, p.D482G, and p.N591S [39]. To elucidate the mechanisms of the disease, knockout rodent models have been developed, however, they fail to accurately recapitulate PFIC2 phenotypes due to the species-specific differences in liver transporter function [40,41]. Alternative efforts to reproduce the disease phenotype *in vitro* using ABCB11-overexpressing non-liver cell lines, hepatocellular carcinoma cells, or PHH from patients with PFIC2 have equally proved ineffective due to the lack of physiological hepatic function (non-liver cell lines), improper protein localisation (hepatocellular carcinoma cells), or donor scarcity (patient PHH) [42]. To overcome these limitations, recent studies have successfully described the generation of two heterozygous PFIC2 hepatocyte models from patient stem cells harbouring the c.−24C > A/c.2417G > A (p.G806D) and c.2782C>T (R928X)/c.3268 C>T (R1090X) mutations; the studies revealed absence of transporter expression in the hepatocyte membrane alongside defects in the apical export of bile acids, highlighting the advantages of iPSC-derived hepatocyte models for the study of disease pathophysiology and novel therapeutic screening [43,44]. Expanding this data set, we now describe, for the first time, the development of a homozygous CRISPR/Cas9-derived PFIC2 model from iPSCs harbouring the common missense mutation D482G, showing significant differences in ABCB11 expression and membrane localisation, in addition to impaired bile acid transport towards the apical side of the cell system. Crucially, we did not observe any compromise in monolayer integrity as confirmed by FITC-Dextran permeability assays (data not shown).

To conclude, we have demonstrated the successful generation of highly functional iPSC-derived hepatocytes. These cells do not only exhibit key hepatic functions, but also provide a valuable model for studying liver biology, disease mechanisms, and potential therapeutic interventions. The detailed cell characterisation presented in this study lays the groundwork for their application in drug discovery, toxicology studies, and regenerative medicine.

## Author contributions

Conceptualisation, C.G., L.P., N.N.; Methodology, C.G., L.P., N.N.; Investigation, M.K., G.G., S.C., G.K., C.R., I.K.; Writing - Original draft, M.L., G.G., S.C., G.K., N.N.; Writing - Review & Editing, M.L., G.G., S.C., G.K., N.N.; Supervision, C.G., L.P., N.N.

## Disclosure Summary

Nothing to declare.

## Acknowledgments

This work was supported by DefiniGEN Ltd.

## Supplementary figure legends

**Suppl. figure 1: (A)** mRNA expression levels of Phosphoenolpyruvate carboxykinase 1 (*PCK1)* in HepG2 cells, wild type Opti-HEP, and primary human hepatocytes (PHH). **(B)** mRNA expression levels of *PCK1* following stimulation with the gluconeogenesis inducer dibutyryl-cAMP (dbcAMP; 0.1 mM) in HepG2 cells and wild type Opti-HEP. Data presented as mean+SEM of 3-4 independent experiments.

**Suppl. figure 2: (A-B)** mRNA expression levels of Phase I CYP450 genes *CYP2A1* and *CYP2E1* in HepG2 cells, wild-type Opti-HEP, and primary human hepatocytes (PHH). mRNA data were normalised to housekeeping gene *18SrRNA* and are presented as mean±SEM of n=2-3 independent experiments. For PHH, cells from three independent donors were used.

**Suppl. figure 3: (A)** mRNA expression levels of key iPSC markers: Octamer-Binding Transcription Factor 4 (*OCT4)*, SRY-Box Transcription Factor 2 (*SOX2)* and Nanog Homeobox (*NANOG*) in wild-type, ornithine transcarbamylase deficiency (OTCD), and Citrullinemia (ASSI) induced pluripotent stem cells (iPSC). **(B-C)** Protein expression of the key iPSC markers OCT4 (green), SOX2 (red), and NANOG (red) in OTCD and ASSI iPSCs. mRNA data were normalised to housekeeping gene Glyceraldehyde-3-Phosphate Dehydrogenase (*GAPDH*) and are presented as mean±SEM of n=2 independent experiments. Nuclei were counterstained with DAPI (blue). Objective: 20x.

**Suppl. figure 4: (A)** Schematic diagram of the D482G mutation in *ABCB11* gene in mutant induced pluripotent stem cells (iPSCs). **(B)** Sanger sequencing chromatogram showing the presence of D482G mutation (GAT > GGT) in PFIC2-mutant iPSCs compared to isogenic control (wild-type iPSCs). The D482G mutation is highlighted in yellow. **(C)** PFIC2-mutant iPSCs were analysed by immunocytochemistry and stained for Octamer-Binding Transcription Factor 4 (OCT4 - green), Nanog homeobox (NANOG - red), and SRY-Box Transcription Factor 2 (SOX2 - red). **(D)** mRNA expression levels of *ABCB11* in monolayer and transwell cultures of wild type Opti-HEP. mRNA data were normalised to housekeeping gene *18SrRNA*, and presented as mean±SEM fold change of n=2-3 independent experiments. Nuclei were counterstained with DAPI (blue). Scale bar: 200 µM.

## Notes

### Competing Interest Statement

The authors have declared no competing interest.

