## Supplementary figures for "Development of optimised human iPSC-derived hepatocytes with improved liver function for *in vitro* metabolic disease modelling and toxicity studies"

### Suppl. figure 1

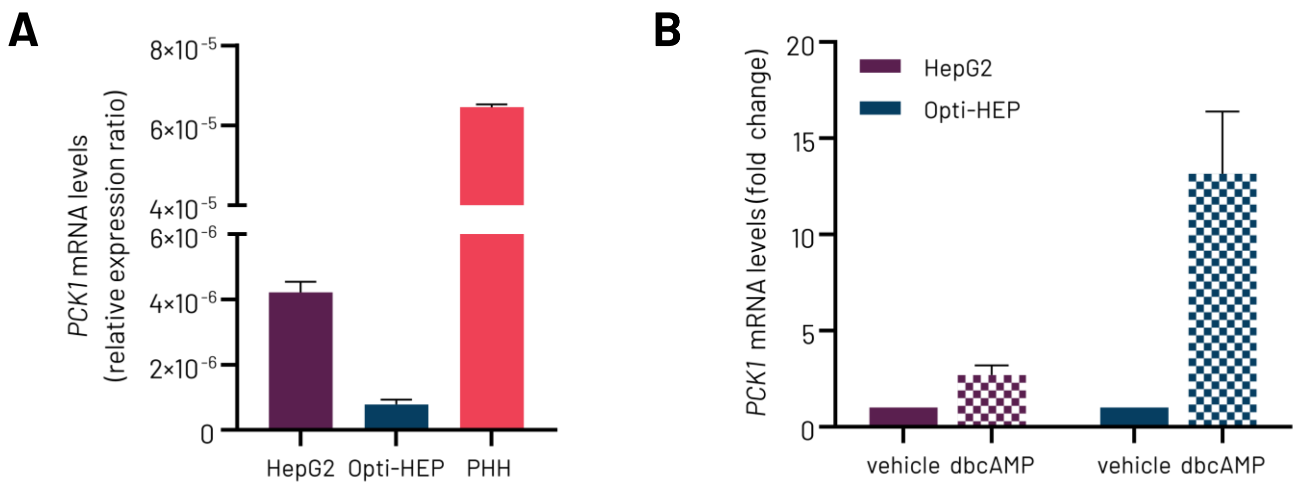

### Suppl. figure 2

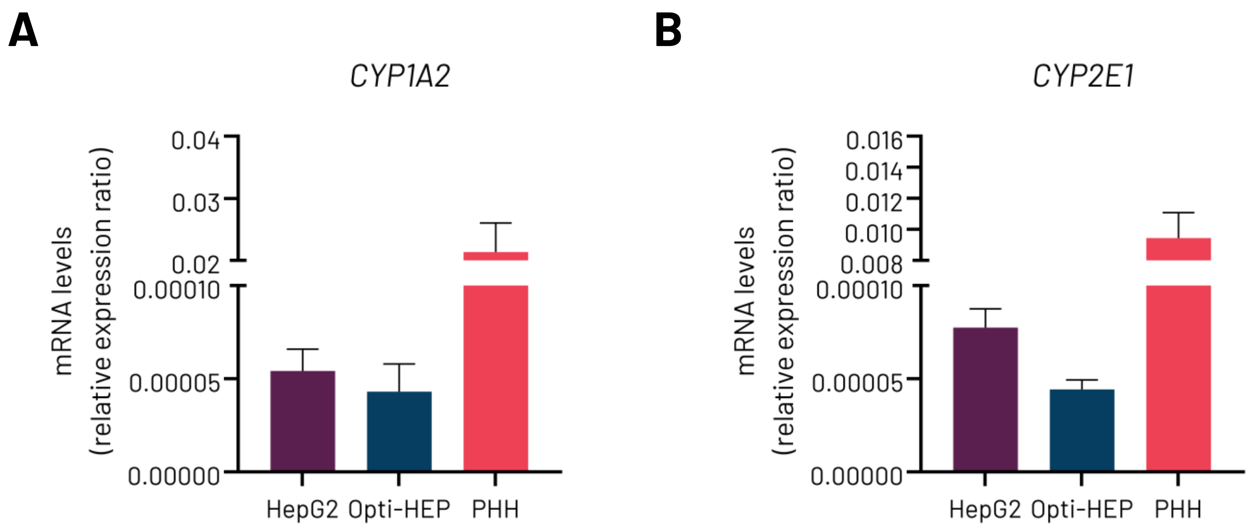

#### Suppl. figure 3

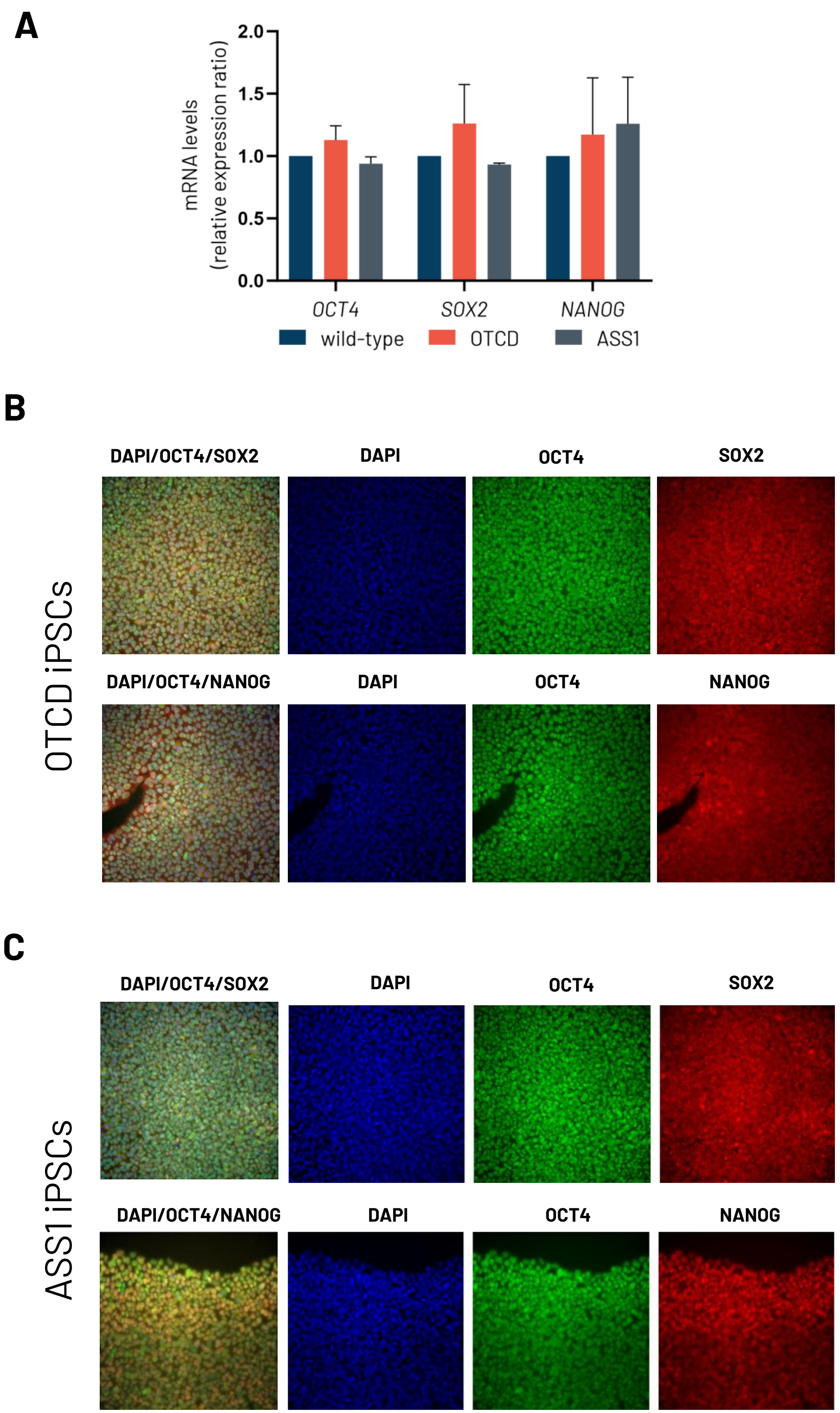

### Suppl. figure 4

**A**

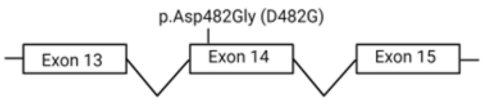


**B**

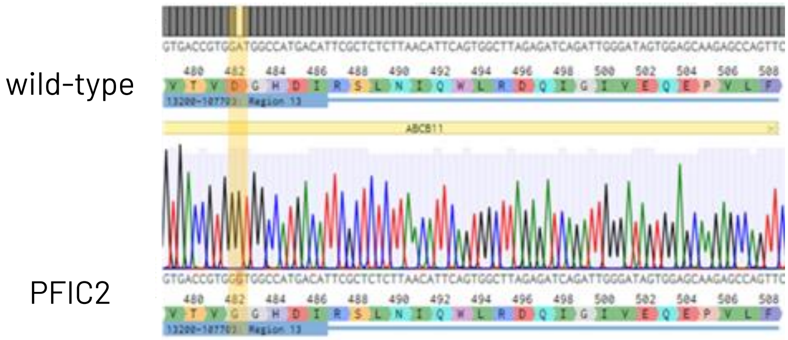


**C**

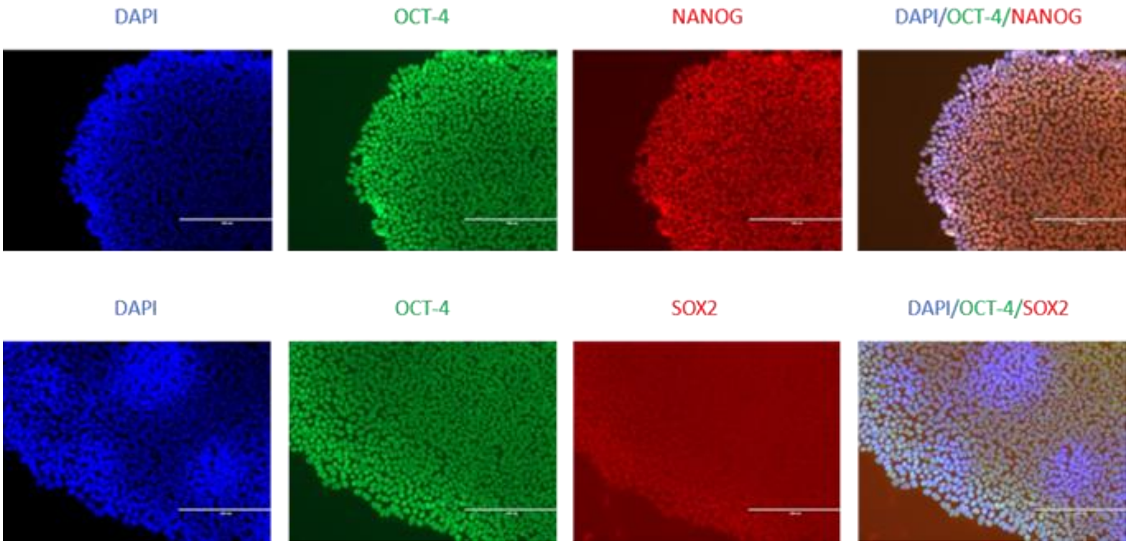


**D**

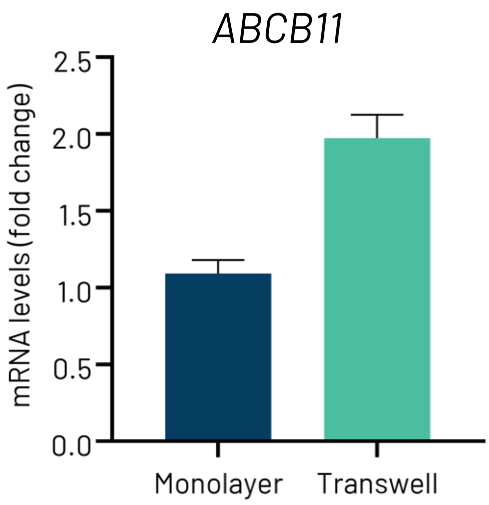

### Suppl. table 1

List of sgRNAs, ssODNs and genotyping primers used for CRISPR-Cas9 gene editing

| Target gene | Mutation | sgRNA Sequence | ssODN Sequence | Forward primer | Reverse primer |
|---|---|---|---|---|---|
| OTC | D175V (Exon 5) | GGAGCGUGAGGUAAUCAGCC | CAATTATCAATGGGCTGTCAGATTTGTACCATCCTATCCAGATTCTGGCTGTTTACCTCACGCTCCAGGTTGGTTTATTTATTTGTCTTACAAAAGAGCA | GGTCACTGGATAAGTCGTGGG | GTCTTCACTCCCAAGCCACA |
| ASS1 | G390R (Exon 14) | UCCCUCAGGUGAGAAGCUCA | CTCCTTGCAGCATGAACGTGCAGGGTGATTATGAGCCAACTGATGCCACCAGGTTCATCAACATCAATTCCCTCAGGTGAGAAGCTCAGGACCCTGACGG | TTCAATGGAAAGCCGACGTG | GAACCTAGGACCCCAATGCC |
| ABCB11 | D482G (Exon 14) | GCGAAUGUCAUGGCCAUCCA | AAGATTCCAAAAGTTGTGATGTTGTGCCCATGCTCTCTAGGTGACAGTGGGTGGCCATGACATTCGCTCTCTTAACATTCAGTGGCTTAGAGATCAGATTG | ACTGTCTCTACTCTTTACTGGAATC | TGACTTGGGAATCATACGAGAAG |

### Suppl. table 2

List of TaqMan probes used for qPCR

| Target gene | Cat.No. | Dye | Probe ID |
|---|---|---|---|
| *GAPDH* | 4326317E | VIC-MGB | Hs99999905_m1 |
| *18S* | 4319413E | VIC-MGB | Hs99999901_s1 |
| *PPIA* | 4326316E | VIC-MGB | Hs99999904_m1 |
| *OCT4* | 4331182 | FAM-MGB | Hs00999634_gH |
| *NANOG* | 4331182 | FAM-MGB | Hs04399610_g1 |
| *SOX2* | 4331182 | FAM-MGB | Hs01053049_s1 |
| *ALB* | 4331182 | FAM-MGB | Hs00910225_m1 |
| *A1AT* | 4331182 | FAM-MGB | Hs01097800_m1 |
| *HNF4A* | 4331182 | FAM-MGB | Hs00230853_m1 |
| *CYP3A4* | 4331182 | FAM-MGB | Hs00604506_m1 |
| *CY2B6* | 4331182 | FAM-MGB | Hs04183483_g1 |
| *CYP2C9* | 4331182 | FAM-MGB | Hs02383631_s1 |
| *CYP2C19* | 4331182 | FAM-MGB | Hs04401150_m1 |
| *CYP2A6* | 4331182 | FAM-MGB | Hs00868409_s1 |
| *CYP2J2* | 4331182 | FAM-MGB | Hs00356035_m1 |
| *CYP1A2* | 4331182 | FAM-MGB | Hs00167927_m1 |
| *CYP2E1* | 4331182 | FAM-MGB | Hs00559367_m1 |
| *ABCB11* | 4331182 | FAM-MGB | Hs00994811_m1 |
| *ABCC2* | 4331182 | FAM-MGB | Hs00960489_m1 |
| *ABCG2* | 4331182 | FAM-MGB | Hs01053790_m1 |
| *G6PC* | 4331182 | FAM-MGB | Hs02802676_m1 |
| *PCK1* | 4331182 | FAM-MGB | Hs00159918_m1 |
| *ASGR1* | 4453320 | FAM-MGB | Hs01005019_m1 |
| *GAPDH* | 4333764T | FAM-MGB | Hs99999905_m1 |

### Suppl. table 3

List of antibodies used for immunocytochemistry

| Target (Dye) | Species | Supplier | Cat. No. | Dilution |
|---|---|---|---|---|
| **Primary antibodies** | | | | |
| OCT4 | Mouse | Santa Cruz Biotechnology | Sc5279 | 1:100 |
| SOX2 | Rabbit | Abcam | Ab5603 | 1:100 |
| NANOG | Rabbit | Cell Signaling Technology | 4903S | 1:100 |
| HNF4A | Mouse | Abcam | Ab41898 | 1:200 |
| ALB | Rabbit | Abcam | Ab2406 | 1:200 |
| A1AT | Mouse | Santa Cruz Biotechnology | sc-59438 | 1:100 |
| AFP | Rabbit | Agilent | A000829-2 | 1:100 |
| ASGR1 | Mouse | BDPharmingen | 563654 | 1:200 |
| E-cadherin | Rabbit | Abcam | Ab40772 | 1:500 |
| ABCB11 | Rabbit | Sigma | HPA019035 | 1:250 |
| MRP2 | Mouse | Abcam | Ab3373 | 1:100 |
| **Secondary antibodies** | | | | |
| Anti-rabbit (AF488) | Donkey | ThermoFisher | A21206 | 1:1000 |
| Anti-rabbit (AF568) | Donkey | ThermoFisher | A10042 | 1:1000 |
| Anti-mouse (AF488) | Donkey | ThermoFisher | A21202 | 1:1000 |
| Anti-mouse (AF568) | Donkey | ThermoFisher | A10037 | 1:1000 |
| Phalloidin (AF488) | | ThermoFisher | A12379 | 1:200 |

### Suppl. table 4

List of antibodies used for western blotting

| Target | Species | Supplier | Cat. No. | Dilution |
|---|---|---|---|---|
| OTC | mouse | Abnova | MAB15702 | 1:500 |
| ASS1 | mouse | Santa Cruz Biotechnology | sc-365475 | 1:100 |
| ASL | mouse | Santa Cruz Biotechnology | sc-374353 | 1:100 |
| CPS1 | mouse | Santa Cruz Biotechnology | sc-376190 | 1:100 |
| ARG1 | mouse | Santa Cruz Biotechnology | sc-166920 | 1:100 |
| ABCB11 | mouse | Santa Cruz Biotechnology | sc-74500 | 1:100 |
| β-actin | rabbit | Cell Signaling Technology | 4970S | 1:1000 |

### Suppl. table 5

List of compounds used for predictive DILI

| Compound | DILI severity category | Cmax Total (μM) | Cmax reference |
|---|---|---|---|
| Ambrisentan | 5 (No liver injury) | 3.03 | Kenna et al 2015 |
| Benzbromarone | 1 (Severe DILI: withdrawn or black box warning due to hepatotoxicity) | 23.58 | Regenthal et al 1999 |
| Clozapine | 2 (High clinical DILI concern: evidence of acute liver failure) | 2.45 | Schulz & Schmold 2003 |
| Fluoxetine | 3 (Low clinical DILI concern: symptomatic liver injury, but not acute failure) | 0.05 | Williams et al 2019 |
| Paroxetine | 2 (High clinical DILI concern: evidence of acute liver failure) | 0.20 | O'Brien et al 2006 |
| Troglitazone | 1 (Severe DILI: withdrawn or black box warning due to hepatotoxicity) | 6.39 | Xu et al 2008 |
| Zomepirac | 5 (No liver injury) | 13.71 | Regenthal et al 1999 |
